# Nucleosome induced homology recognition in chromatin

**DOI:** 10.1101/2021.04.29.441844

**Authors:** Jonathan G. Hedley, Vladimir B. Teif, Alexei A. Kornyshev

## Abstract

One of the least understood properties of chromatin is the ability of its similar regions to recognise each other through weak interactions. Theories based on electrostatic interactions between helical macromolecules suggest that the ability to recognize sequence homology is an innate property of the non-ideal helical structure of DNA. However, this theory does not account for nucleosomal packing of DNA. Can homologous DNA sequences recognize each other while wrapped up in the nucleosomes? Can structural homology arise at the level of nucleosome arrays? Here we present a theoretical investigation of the recognition-potential-well between chromatin fibers sliding against each other. This well is different to the one predicted and observed for bare DNA; the minima in energy do not correspond to literal juxtaposition, but are shifted by approximately half the nucleosome repeat length. The presence of this potential-well suggests that nucleosome positioning may induce mutual sequence recognition between chromatin fibers and facilitate formation of chromatin nanodomains. This has implications for nucleosome arrays enclosed between CTCF-cohesin boundaries, which may form stiffer stem-like structures instead of flexible entropically favourable loops. We also consider switches between chromatin states, e.g., through acetylation/deacetylation of histones, and discuss nucleosome-induced recognition as a precursory stage of genetic recombination.

## I. INTRODUCTION

**K**ornyshev-**L**eikin (KL) theory and its extensions^1–4^, as well as other theories of DNA recognition^5,6^, suggest that homologous tracts of two different DNA molecules may associate with each other due to the physical interactions between them alone, in the absence of proteins. This effect was confirmed *by Inoue et al*. in 2007^7^ and has been further demonstrated in a series of *in vitro* experiments, including liquid crystalline ordering in spherulites^8^ and magnetic-bead single-molecule techniques^9,10^. These results all indicate a preferential interaction of homologous sequences over non-homologous sequences, which is enhanced by the effects of osmotic pressure, crowding agents and confinement. While these models are applicable to bare DNA molecules for simple biological systems, once we begin to consider eukaryotic systems, we need to account for the packing of DNA in chromatin structures with the help of nucleosomes.

The current view of chromatin is that of a liquid/polymer system of irregularly (but not randomly) packed beads-on-the-string nucleosome arrays, sometimes referred to as the 10-nm fiber^11–13^. While mostly amorphous, the organisation of nucleosome arrays is characterised by microdomains with distinct properties, sometimes characterised by local alignment^14,15^. In many cases, local alignments of nucleosome arrays are directed by homotypic DNA sequence repeats such as L1 and B1/Alu^16^.The theoretical foundations of such interactions are currently not well established – this is a problem we aim to address in this paper.

At a larger scale, electrostatic repulsion and entropic contributions are expected to push the nucleosome arrays apart from each other, which is counteracted by osmotic stress and confinement in the cell nucleus^17,18^ and further modulated by the composition of hydrophobic/hydrophilic residues^19^. On top of these generic forces act DNA sequence-specific bridges formed by proteins. The most prominent example is the demarcation of the genome by the architectural protein CTCF, which instructs its partner, the ring-shaped molecular motor Cohesin, where to form a DNA-DNA bridge^20,21^. Between such bridges lie chromatin loops, which are usually depicted as loosely constrained. However, if there does exist some internal recognition of the structural homology of nucleosome arrays, it could potentially affect the structure of these loops. This work seeks to investigate if this effect is present and significant, and its potential implications.

Another potential effect of the structural homology of nucleosome arrays is its modulation of the ability of DNA to mutually recognise homologous sequences in processes like homologous recombination. The initial alignment of the two DNA tracts minimises potential mistakes in gene shuffling between two parental copies of DNA in meiosis and DNA repair. Following double strand breaks in DNA, RecA proteins promote the association of homologous DNA duplexes through the formation of DNA-RecA filaments^22^. The initial alignment of the two DNA tracts minimises potential mistakes in gene shuffling between two parental copies of DNA in meiosis and DNA repair. Following double strand breaks in DNA, RecA proteins promote the association of homologous DNA duplexes through the formation of DNA-RecA filaments. The initial sequence probing only requires eight base pairs (bp), which enables the elimination of a large proportion of mismatched sequences within the genome. However, within the human genome, any given 8 bp sequence can be encountered multiple times. Therefore, this initial rapid testing is then followed by slower sequence testing in later stages of the recombination reaction, aided by ATP hydrolysis and partly controlled by RecA and other proteins. This ‘proofreading’ process minimizes the chances for erroneous recombination, but the process will be very much sped up by the initial juxtaposition of the right sequences. While the recognition of filaments composed of DNA coated by RecA is different from the structural recognition of nucleosome arrays, experimental studies have shown that homologous DNA molecules also preferentially associate with one another in the absence of protein helpers. This indicates that energy dependent mechanisms controlled by molecular motors can be complemented by ‘coarse-grained’ sequence recognition mechanisms, which may work in cooperation to improve the accuracy of this process. For comparison, in bacterial recombination, homology of 50-200 bp is required^23–25^. In eukaryotic systems, DNA is hidden by histones, and therefore recognition may occur as the result of a two-step process, where we initially observe a ‘coarse-grained’ alignment of similarly structured nucleosome arrays, followed by a more precise DNA-level match once initial alignment has been made.

To investigate the effect of structural homology of nucleosome arrays on the processes mentioned above, one has to consider partial or full unfolding of the nucleosome that allows the juxtaposition of naked DNA sequences. Temporary nucleosome unwrapping and repositioning does indeed happen *in vivo*, facilitated by thermal fluctuations^26^ and active energy-dependent chromatin remodelling^27^. However, histones bind tightly to DNA with an energy on the order of ∼20-30 *k*B*T*^28^, and so it is energetically costly to fully remove them from these regions. It is therefore not unreasonable to expect that nucleosomes play some role in the sequence recognition between homologous DNAs. Indications on this were noted long before DNA was discovered as bearer of genetic information [c.f. McClintock^29^, ‘*there is a tendency for chromosomes to associate 2-by-2 in the prophase of meiosis*’]. Additionally, *in vitro* experiments of short DNA sequences with histones also show evidence of preferential association of identical sequences^30^. However, despite these observations, there also exists experimental evidence that an increased nucleosome density decreases the efficiency of homologous recombination^31–33^. We discuss these results alongside the theory presented here later in this paper.

## II BASIC APPROACH AND THE MODEL

Here, in the spirit of the KL-theory^1–4^, but using its simplified, achiral implementation, we model the effect of nucleosome arrangement on the structural homology recognition between nucleosome arrays. If we consider two long rod-like molecules (*ν* = 1,2) with parallel, cylindrical, water-impermeable cores, we can derive the interaction energy per unit length using a mean-field formalism within the Debye-Bjerrum approximation:

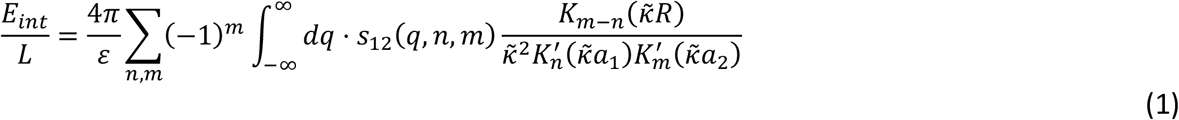

where *ε* is the dielectric constant of the medium (for water, *ε* ≈ 80), *K*_*n*_(*x*) is the n^th^ order modified Bessel Function of the second kind and 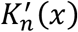 is its derivative with respect to *x, R* is the interaxial separation between the two molecules, *a*_1_ and *a*_2_ are the radii of the two molecules, 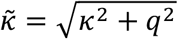, where *κ* is the inverse Debye length, and *s*_12_(*q, n, m*) is the charge density correlation function, given by:

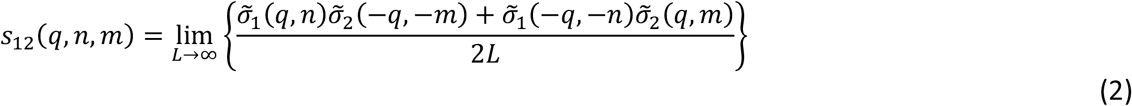

where 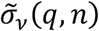 is the Fourier transform of the cylindrical surface charge density of the interacting nucleosome arrays, the subscripts *ν* = 1,2 label the two interacting fibers, and *L* is their length. Through these equations, we can link the structure of the charge distributions on the molecules to the electrostatic interaction between them. Such an approach allows calculation of the recognition energy, that is, the preferential interaction energy between juxtaposed homologous and non-homologous sequences^3^.

To use this approach, we need to consider the surface charge distribution of chromatin, and hence its structure. Positively charged histones partially compensate the strongly negatively charged DNA backbone, which facilitates condensation into a number of higher order structures with the help of architectural proteins such as CTCF and cohesin. Since the chromatin fibers have similar structures, we can construct a simple surface charge model of chromatin which can describe both systems. From this, we can obtain the relevant interaction energies and hence investigate the ‘histone-on’ homology recognition well, and its implications for these biological effects. A possibility of recognition of structurally homologous chromatin fibers has previously been considered by *Cherstvy & Teif*^34^. This study led to the conclusion that long range recognition is facilitated by protein bridging interactions, and when closely juxtaposed, direct electrostatic recognition enables fine-tuned structure-specific recognition. In the present paper we explore an opportunity for the fibers with nucleosomes to recognize their homology in a ‘collective’ manner, but without protein bridging interactions. We will explore the conditions for forming a ‘histone-on’ homology recognition well, investigate the shape of that well, and the possible implementations of these findings for the biological processes mentioned above.

Here we take a crude, coarse-grained approach to the structure of chromatin, where we describe the nucleosomes as discrete cylindrical blocks of charge distributed along a negatively charged cylindrical core representing the linker DNA, as shown in Figure 1. While this does neglect the helicity of the linker DNA, the inclusion of nucleosomes restricts how close the DNA in the chromatin fibers can come to each other. The combination of this restricted proximity and the strong electric screening (Debye length *λ*_*D*_ ≈ 7 Å in physiological conditions) allows us to make the assumption that the helicity will most likely have a moderate effect on the gene-gene interactions in the ‘histone-on’ state. Additionally, the Debye-Bjerrum approximation under which the interaction energy was derived may be not a bad first approximation towards description of intermediate to large separations between molecules; at these length scales the direct DNA-DNA contribution will be less important.

**Figure 1:**
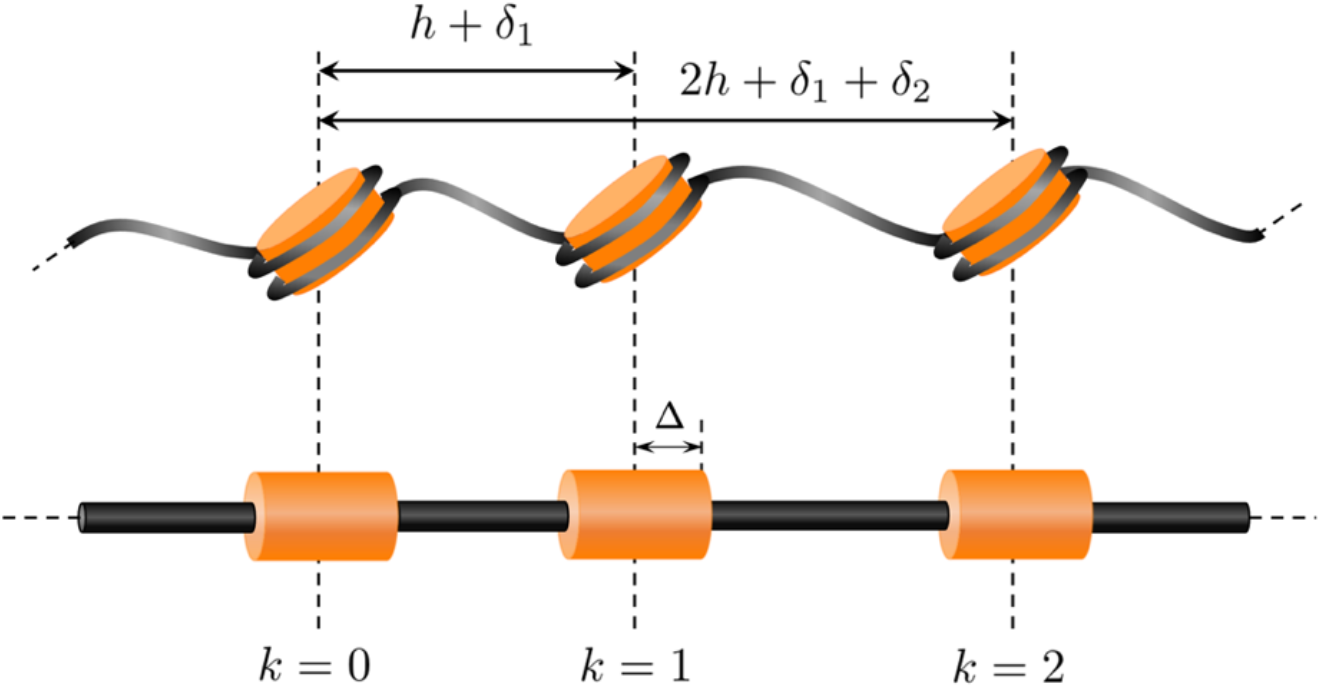
A diagram to show the model used for the 10-nm chromatin fibre in this study. The orange cylinders represent the nucleosomes, with half-width △ = 5 nm and radius r_nuc_ = 5 nm. Their centres are distributed according to an accumulated disorder, where the random variable 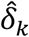 is defined as the fuzziness, relating to the disorder in nucleosome positioning. The black cylinders represent the linker DNA, with radius r_DNA_ = 1 nm. The fine details of the helicity of the linker DNA, and the DNA wrapped around the histones are neglected in the model, a reasonable approximation at the length scales in the system.

To account for the difference in radii between nucleosomes and linker DNA, we express the total charge density as a sum of the nucleosome surface charge density 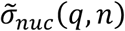 at radius *r*_*nuc*_ and the DNA surface charge density 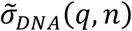at radius *r*_*DNA*_:

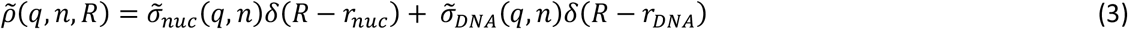

where *δ*(*x*) is the Dirac delta function, and 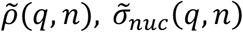 and 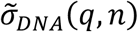 are the Fourier transforms of the real space cylindrical charge densities:

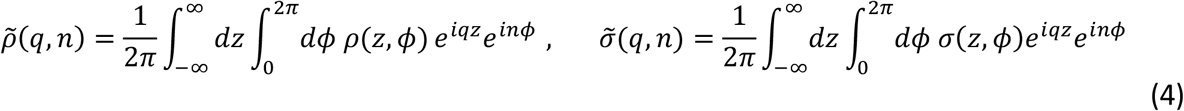

As justified above, we express the DNA surface charge density as a continuous line charge of length *L* centred around zero, with charge density 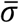. For the nucleosomes, we sum rectangle functions centred around *z*_*k*_ with half-width Δ, such that:

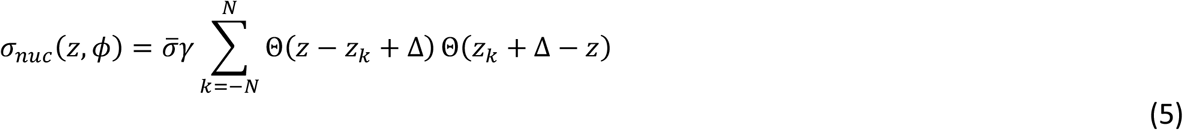

where Θ(*z*) is the Heaviside step function, and *γ* is a tuneable parameter representing the charge compensation of the DNA by the histone, such that 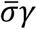 is the net charge density on the nucleosome. This expression leads us to discuss how to model the nucleosome distribution through their centres, *z*_*k*_. Having centred the two molecules around *z* = 0, we can initially distribute the nucleosomes periodically, such that *z*_*k*_ = *kh*, where *h* is the average nucleosome repeat length (NRL). However, we do expect a deviation from ideal periodicity in this distribution. Nucleosome positioning is a dynamic process, but for simplicity we approach the distribution on a time-averaged basis. As we are investigating the innate structure of chromatin in this work, we can also neglect the effect of nucleosome remodelling enzymes and transcription factors, which affect the positioning of nucleosomes. Nucleosomes have preferential positions on DNA^35,36^, and hence two homologous DNA tracts are expected to have similar nucleosomal positioning, correlated with their sequences. In particular, Hi-C data has suggested inter-chromosomal co-localisation of Short Interspersed Nuclear Elements (SINEs), including Alu sequences^37^. Each Alu sequence usually positions 1-2 nucleosomes^38^. However, since each well-positioned nucleosome or well-located nucleosome-depletion provides a boundary arranging about 10 other nucleosomes^39^, such DNA sequences provide long-range nucleosome organisation effects. We will see in a moment how this can be taken into account.

A naïve approach would be to consider *long-range* order, where nucleosome centres are distributed as 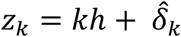 with 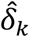 standing for random displacements of the *k*^,-^ nucleosome. This random variable 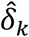 can be described as the *fuzziness* of the nucleosome positions, as it indicates the degree of disorder in nucleosome centres. Here, the periodicity of the nucleosomes persists along the length of the molecule, and fluctuations in nucleosome position are described by a kind of ‘static Debye-Waller smearing’ of the lattice. However, this does not accurately describe deviations from ideality in DNA and chromatin. Indeed, the preferential positions for nucleosomes are correlated with the nucleotide sequences; thus, on average, identical/similar sequences are expected to have the nucleosomes at the same positions within the sequence. Therefore, a better choice for the nucleosome distribution would be to follow the model of non-ideality in the DNA double helical structure; that is, an accumulation of disorder in twist angle Φ(*z*)^40^. Interestingly, even before the structure of DNA was deciphered, Schrödinger, in a way, acknowledged the key distinction between these long-range and short-range order models in his take on ‘*What is life?’*, when he hypothesised that the genetic material must be some sort of ‘aperiodic crystal’^41^. In our model, this aperiodicity or accumulation of disorder plays a key role in the ability for DNA and chromatin to recognise each other, distinguishing the cases when the disorder is identically accumulated, and when it is not correlated, between the molecules. All in all, if DNA sequence defines some preferential nucleosome positioning, two homologous genes may have nucleosomes in irregular, but overall similar arrangements, whereas in the case of two nonhomologous genes, the arrangements may be entirely different – random with respect to each other. As we will show below, this circumstance may let the genes recognise DNA sequence homology at the structural level. We hence define our nucleosome centres, *z*_*k*_, using an accumulated disorder^42^, according to this short-range order model (*see Figure 1*):

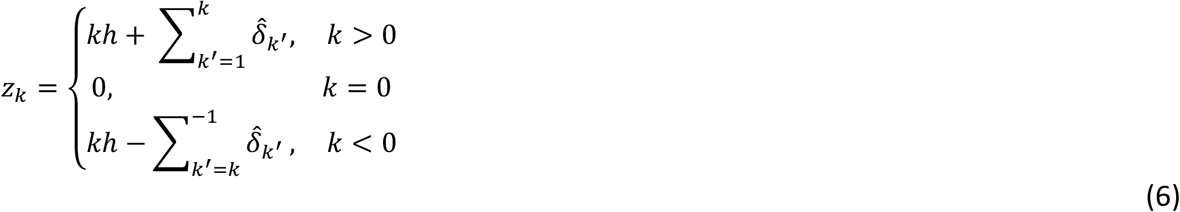

In the helical theory, it was found that disorder propagates from the centre of the molecule by minimising the interaction energy with respect to *k*, and so we set *z*_*k*_ = 0 when *k* = 0. The introduction of these fluctuations allows us to model the interaction of both homologous and non-homologous molecules. For homologous DNA texts, we can assume that the disorder on the two DNAs ‘accumulates’ in the same way, i.e., 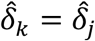., for *k* = *j*, whereas for non-homologous DNA texts, we assume that 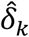 and 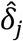 are uncorrelated, so that for long tracts there will be no similarity in nucleosome positioning along their lengths.

This assumption of an identical thermodynamically favoured distribution of histones on DNA molecules of the same base-pair sequences, sometimes referred to as the nucleosome positioning code, has a number of supporting evidences^43,44^. It is known, however, that in the human genome only ∼38% of the nucleosome occupancy *in vivo* is stably encoded in the DNA sequence^45^. In addition, some degeneracy in the nucleosome positioning may disturb the commensurability of the related charge distributions on homologous molecules. If, for a moment, we step away from chromatin and recall the electrostatic recognition mechanism between bare DNA molecules, we must note that homologous DNA sequences, *do not* have absolutely identical sequences of base pairs, identical structure, and hence they do not have identical charge distributions. They are, however, *almost identical*, and our earlier analysis has shown that the rare local deviations from the identity of homologous sequences have a negligible effect on the interactions between homologues^46^. This is because the presence of such deviations does not have an accumulating effect (which lies in the heart of this recognition mechanism). Similarly, we expect only a minor distraction of commensurability between the charge distributions of histones on opposing homologous sequences. But again, the distribution of nucleosomes, even on identical sequences, may not necessarily be identical. Moving a nucleosome out of its preferential position may require 20-30 kBT of energy^28^ and the action of molecular motors – chromatin remodellers^27^. Thus, the assumption of similar nucleosome-related charge distributions for homologous DNAs may be a reasonable approximation for a large fraction of chromatin.

It is important to note that this ‘structural homology’ of chromatin fibres is not necessarily only dependent on DNA sequence homology. A simple boundary along the chromatin fibre, such as a CTCF protein, would arrange neighbouring nucleosomes in a regular array. Thus, the model for nucleosome positioning formulated here is applicable to both the case where the nucleosome distribution is dictated by DNA sequence, and the case where it can be dictated simply by boundary conditions (or any other effects). We discuss results for both applications of the model in section III.

### HOMOLOGY RECOGNITION OF SLIDING CHROMATIN MOLECULES CAN ALIGN THEM IN FAVORABLE JUXTAPOSITION

Using the basic formulae of the theory, and the crude coarse-grained model constructed above, we calculate the recognition energy profile for the sliding of one long chromatin fiber against another within a juxtaposition window of length *L*_*J*_, as described schematically in Figure 2. We determine the recognition well by finding the difference between energies of two homologous chromatin fibers sliding against each other and two non-homologous fibers sliding against each other. In the juxtaposition window, that is, the region over which the interaction is calculated, we treat the fibers as straight and rigid. The inclusion of torsional flexibility would reduce the overall recognition energy^47^ and so the system we describe here would yield the deepest well possible. This approach is more realistic in the case of homologous recombination, as it does not require the entire length of each fiber to be in juxtaposition with the other. Instead, the fibers only need to lie closely together within a defined length of the order of the persistence length of the fiber, or larger if the genetic machinery within the cell (which may control the sliding of the fibers), can provide a longer juxtaposition. There is no clear consensus on the persistence length of nucleosome arrays, but available estimates from yeast studies suggest the value similar to the one for bare DNA, namely around 50 nm, with compaction ∼50bp/nm^48,49^ (which is on the order of ∼10-15 nucleosomes). The overall total length of the fibers is considered to be significantly longer than the juxtaposition window, and so we can neglect potential ‘edge effects’.

**Figure 2:**
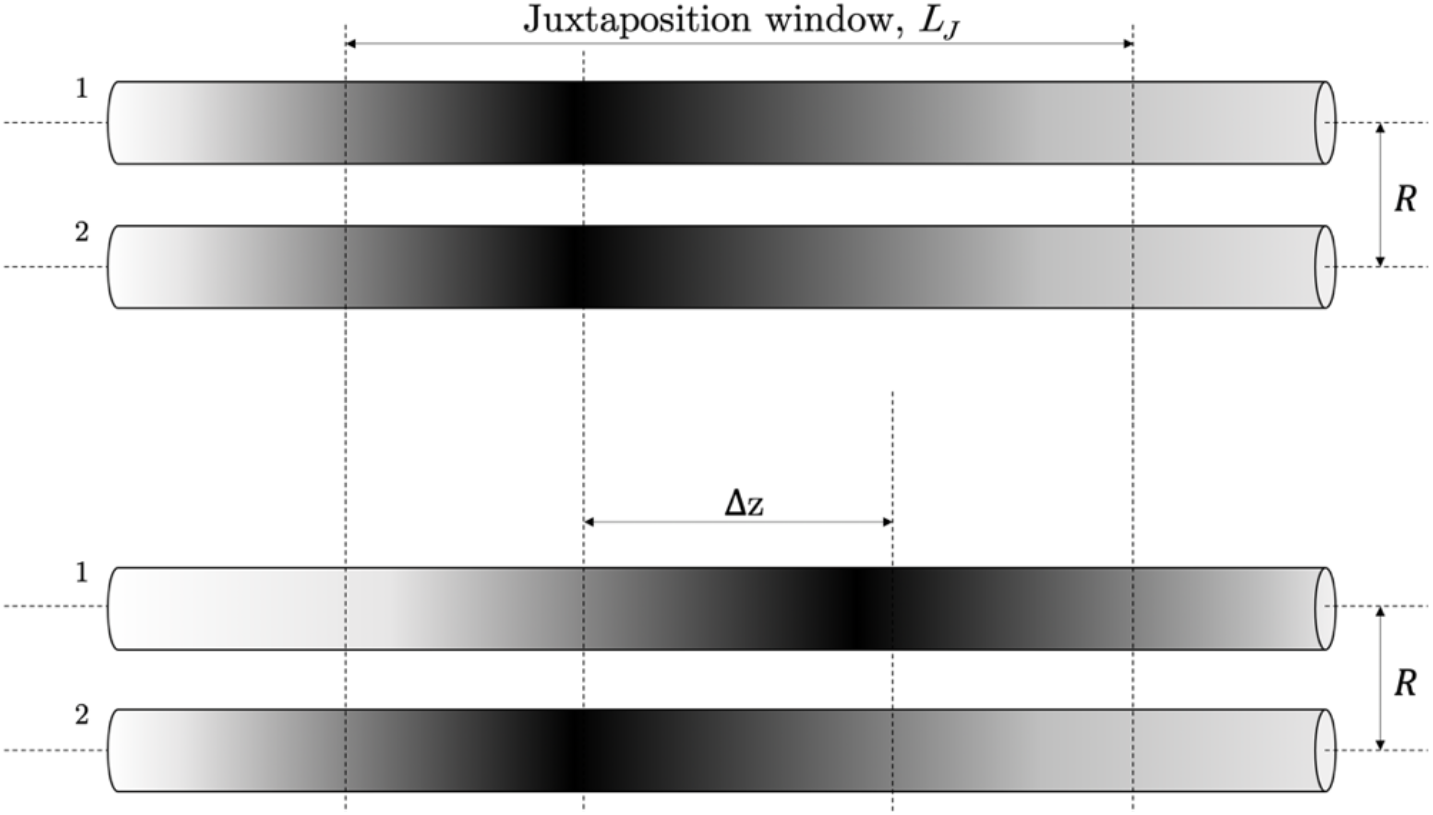
A schematic representation of two long nucleosome arrays interacting within a juxtaposition window of length L_J_. Outside the window, the interaction is neglected (i.e. the molecules lie in close proximity only over the length L_J_). The upper panel shows two homologous sections fully juxtaposed; in the lower panel, array 1 is shifted by a distance |△z|. Homology here is represented by the shading.

Measuring the axial shift as Δ*z* within this framework, and taking a Gaussian average over the random fluctuations, the recognition well is given by:

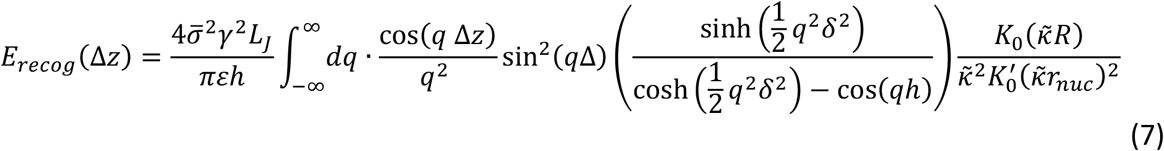

where *r*_*nuc*_ is the radius of the chromatin fiber, and 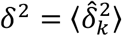, the mean squared fuzziness. The derivation of this formula is presented in the Appendix. A consequence of this formula is that the recognition well depends only on the nucleosome parameters, hence stating that recognition in this coarse-grained model is controlled by the positions and charge of the nucleosomes. This is expected, as we have formulated the model in such a way that it neglects the KL-helix-specific interactions responsible for the homology recognition well between bare DNA, an assumption that may be reasonable at the length scales that we see in chromatin.

Calculating the integral in the r.h.s. of Eq. (6) numerically, we see a series of maxima and minima decaying with axial shift in Fig. 3. All calculated energies are scaled to *k*_B_*T* by expressing the surface charge density in units of *e*Cnm^-2^, where *e* = 1.609 × 10^−19^ C, and hence we can utilise the Bjerrum length, ℓ_*B*_ = *e*^2^/*εk*_*B*_*T*, which in physiological conditions ≈ 0.7 nm. The analytical forms of features of interest in the recognition well are not clear given the complex structure, however, there are similarities to the helical case we can pull on. The decay in maxima and minima is over a *coherence length, ξ*_*c*_, analogous to the helical case, where the helical coherence length of the interaction is given by *λ*_*c*_ = *h*_*r*_/(ΔΩ)^2^, where *h*_*r*_ is the helical rise between base pairs and ΔΩ is the variation in twist angle. In a similar way, we see that the coherence length in the chromatin fibers depends on *h* and *δ*, where it decreases with increased average fuzziness (*δ*). The positions of the minima, dictated by Eq. (6), are also expected to depend on *h* and *δ*, moving further away with increasing nucleosome spacing and variation. The primary maximum at full juxtaposition (Δ*z* = 0) corresponds, obviously, to full alignment of similar charges. This maximum decreases with increasing *δ*, a result of greater positional variance reducing the repulsion between the fibers.

**Figure 3:**
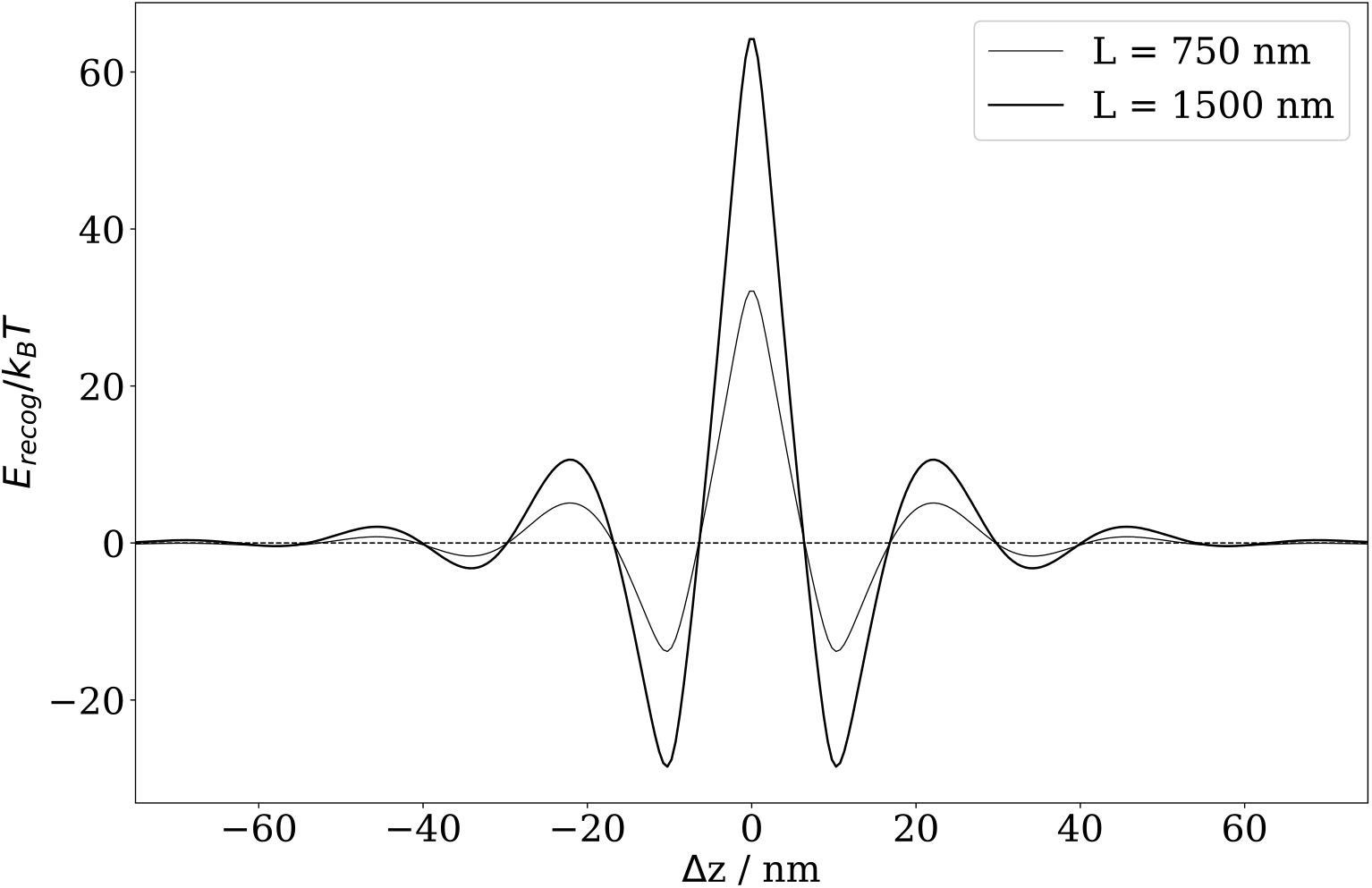
The recognition potential well for the indicated values of juxtaposition length, LJ. The energy is given in units of thermal energy at room temperature. Parameters used in the calculation are: 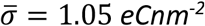, γ = 0.63, κ = 10/7 nm^-1^, Δ = 5 nm, h = 20 nm, R = 11 nm, and r_nuc_ = 5 nm. The average value for δ varies between organisms, but literature generally refers to ‘well-localised’ nucleosomes as having a standard deviation of 10-20 bp (≈3.4 − 6.8 nm)^36^. Here we have used the upper bound δ = 6.8 nm.

The recognition well is also proportional to the juxtaposition window. The larger the window the fibers interact over, the greater the strength of the interaction. There is a problem with this, however. If we seek to minimise the energy, then this proportionality would allow fibers to be trapped in a configuration where they are aligned along their entire length, which in this system is where *L*_*J*_ → ∞. Such a conclusion is of course misleading, as we have not taken into account the entropy of the system. As the length of juxtaposition increases between the two fibers, the number of configurations accessible by the fibers decreases significantly, resulting in a loss of entropy in the system. Physically, the system could perhaps compensate such loss of entropy by the release of waters of hydration associated to each fiber, leading to an overall increase, but this is a pure speculation. Most likely, there will just be a compromise between the energetic and entropic terms, leading to an ideal juxtaposition length, depending on external conditions – crowding agents, confinement, etc. Thus, all we have done so far is the calculation of the interaction energy for typical juxtaposition lengths. At these lengths we can see that the primary minima are at small shifts of Δ*z* ≈ *h*/2 from full alignment and are deep enough to temporarily trap the fibers in this shifted conformation.

As mentioned earlier, ALU sequences and other repetitive elements contribute to the distribution of the nucleosome array, and we can see that observations regarding their association can be predicted by our model. It is important to emphasise however that while one single ALU element, corresponding to two nucleosomes, (*L*_*J*_ ≈ 11 *nm*) can still recognise similar sequences *in vitro*, the ‘electrostatic recognition well’ appears to be too shallow, at least when operating with the macroscopic value of the dielectric constant of water, and the overall recognition effect will be too weak. These results, however, cannot be directly treated with the theory presented here, as the effects that we describe are associated with the juxtaposition of long chromatin fibers.

With that said, we should admit that if the effective dielectric constant at such nanoscale distances appears to be much smaller, the effect can be amplified by an order of magnitude. If the latter is not true, even a weak effect may be sufficient for association *in vitro*, but structural recognition *in vivo* likely arises from several nucleosomes that organised around a single Alu repeat due to the boundary effects exerted by strongly positioned nucleosomes. It is also worth noting that within the real cell environment any sources of recognition would have to compete with many other ‘distracting’ interactions.

### INTERACTION OF FINITE-LENGTH CHROMATIN FRAGMENTS

In many cases it is possible to consider isolated genomic regions of fixed length which have limited interactions with the surrounding genome. One prominent example is the case of nucleosome arrays isolated by CTCF boundaries. The regions between CTCF boundaries sometimes form Topologically Associating Domains (TADs) or sometimes smaller compartments called loops^20,21^. In either case, it is convenient to consider these regions bounded by CTCF as physically independent from the neighbouring regions along the 1D genomic coordinate. In the human or mouse genome, the distance between neighbouring CTCF sites is on average around 10,000 bp^50,51^. For such systems, it may be useful to understand how finite sized nucleosome array fragments interact if they are juxtaposed with each other (Figure 4).

**Figure 4:**
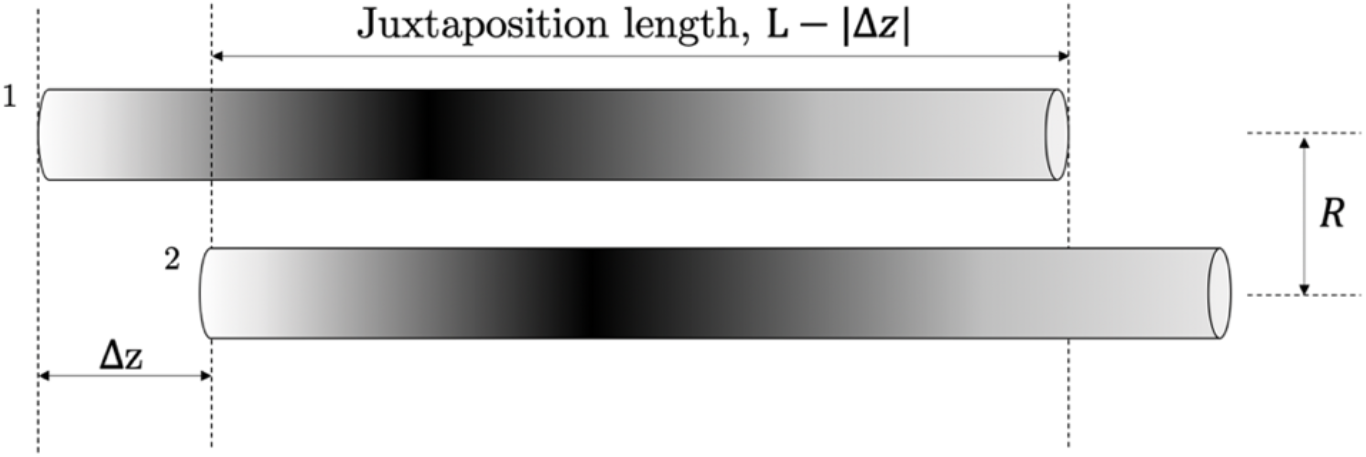
A sketch of a configuration of sliding DNA fragments of finite length L, where the fragments have the same homologous sequence. The juxtaposition length here is L − |Δz|, beyond which no interaction is taken into account.

Let us consider the interaction between finite length chromatin fragments of length *L*. The main difference between this system and the system of infinite nucleosome arrays is that the interaction occurs over a juxtaposition window of *L* − |Δ*z*|, and that the contribution from sections of the molecules not in juxtaposition to the overall interaction may be neglected. Such a situation is sketched in Figure 4. As the interaction energy is proportional to the juxtaposition window length, we expect to see the interaction diminishing proportionally to |Δ*z*|. However, as we saw in the previous system, we expect the recognition to persist over a certain coherence length, *ξ*_*c*_, and past this length, the interaction between homologous and non-homologous nucleosome arrays is identical. Therefore, one may expect little difference in the calculation of the recognition energy from the case considered in the previous section.

The latter is valid to a point: for the system considered in the previous section, the interaction is proportional to *L*_*J*_, whereas in the interaction of finite fragments, the interaction is proportional to *L* − |Δ*z*|. However, as features of interest in the well reside in the region −*ξ*_*c*_ < Δ*z* < *ξ*_*c*_, in the limit where *L* ≫ *ξ*_*c*_, *L* − |Δ*z*| ≈ *L*, for large molecules this expectation would be valid. Still, we wish to carefully investigate the situation with short fragments. The expressions for the interaction energies for this system are presented in the appendix. Their difference yields Eq. (8), the recognition energy for chromatin fragments:

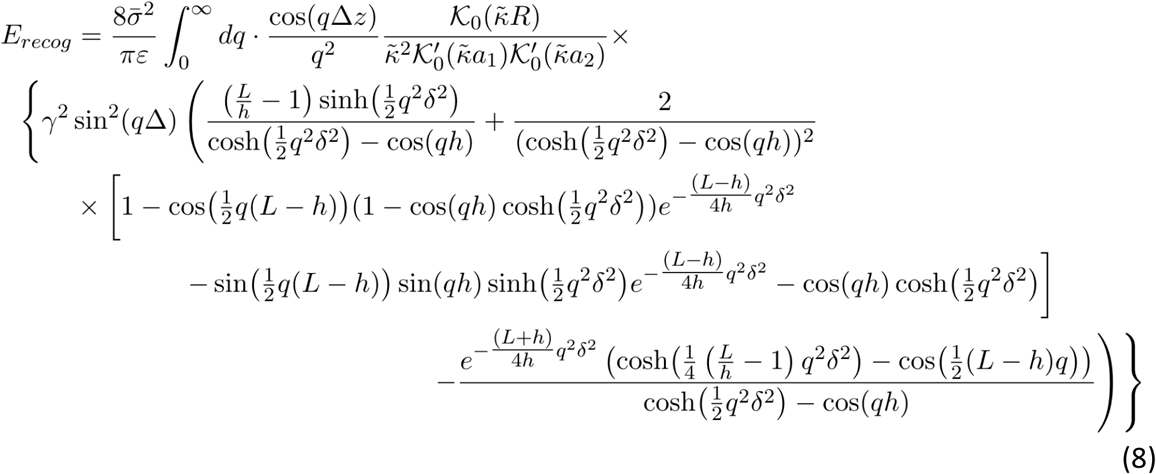

The integrals in Eqs. (8), (A22) and (A23) were numerically calculated and are plotted in Figure 5.

**Figure 5:**
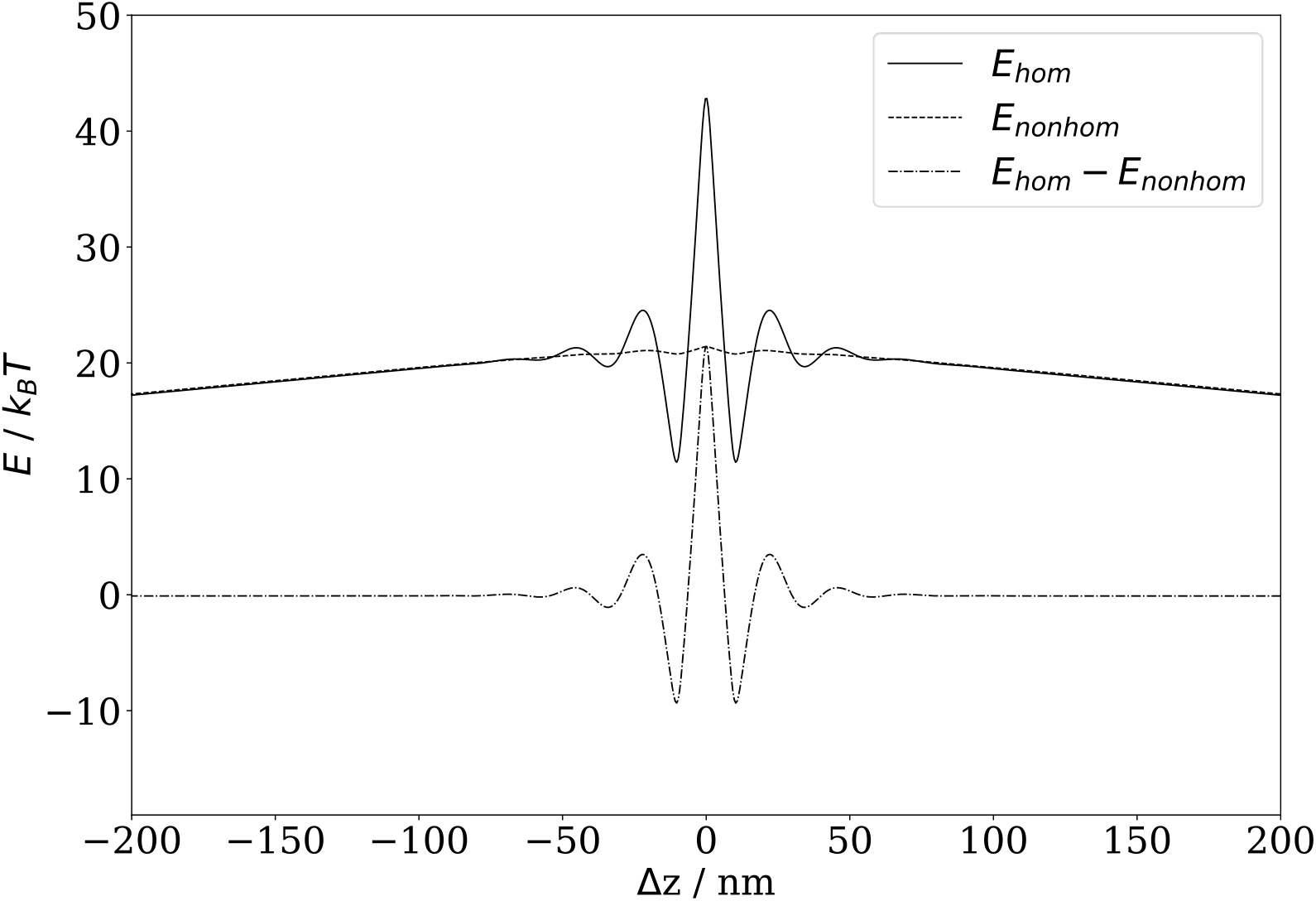
The interaction profiles for homologous and non-homologous distributions of nucleosomes show proportionality to the juxtaposition length, L-|Δz|. Their difference yields the recognition potential well. The homologous energy profile E_hom_ decays to the non-homologous profile E_nonhom_ at large shifts, that is, larger than the coherence length.

The energy profiles for both *E*_*hom*_ and *E*_*n*o*nhom*_ are proportional to *L* − |Δ*z*|. In the non-homologous case, we see no oscillations in the profile as a consequence of the completely uncorrelated deviations in nucleosome centres. This is expected; when averaging over a long fiber, attractive or repulsive interactions in one region of the fiber will be cancelled out by opposite interactions in another region, resulting in a straight-line profile proportional to the juxtaposition length *L* − |Δz|. For homologous nucleosome array fragments, we observe the same series of maxima and minima as in the previous system, which decays to the non-homologous case over the coherence length *ξ*_*c*_. This is also intuitive; when homologous nucleosome arrays are shifted past a certain point, they lose correlation and become equivalent to non-homologous texts. Hence, taking the difference, we obtain a similar recognition well, which converges to the recognition well we have already seen at large *L*, as per the argument laid out above.

The analytical forms of the minima and coherence length have the same dependence on *h* and *δ* that we saw previously. Thus, the finite sizes of interacting nucleosome arrays can be quite well described by the simpler expressions derived in the previous section. It is however still important to develop this finite-sized model for the biologically relevant cases, such as the isolated genomic regions (TADs, loops) detailed above.

Despite the recognition well having similar properties to the previous system, this clarifies that the ‘attractive’ minima do not necessarily indicate direct electrostatic attraction between fibers. Rather, the minima are positions of reduced repulsion; if forced to be in close proximity, for example by protein bridging, and under osmotic stress, the fibers will adopt positions corresponding to these minima.

## III. BIOLOGICAL IMPLICATIONS

Based on the results of the above presented model, we will speculate below about their possible manifestations in molecular genetics.

### Boundary-ordered nucleosome arrays may form stem-structures instead of loopss

Nucleosome arrays are frequently organised with the help of boundaries, such as nucleosome-depleted regions at transcription start sites or binding sites of proteins such as CTCF. A bound CTCF organises up to 20 nucleosomes around it in an ordered array^52,53^. The closer to CTCF, the more ordered this array, and the smaller the distances between neighbouring nucleosomes (Figure 6A)^51^. We previously estimated that in mouse or human cells, up to 10% of the whole genome is organised in such nucleosome arrays with the help of CTCF. Some regions have neighbouring CTCF sites at relatively small distances from each other, enclosing just a few nucleosomes between them (Figure 6B). Other regions have CTCF sites at longer distances, with chromatin fibers between them looping in 3D space. Using the theory developed above, we can ask the following two questions: 1) whether ordered nucleosome arrays near chromatin boundaries interact stronger than ‘fuzzy’ less ordered arrays, and 2) do these interactions provide additional constraints in the chromatin loop structures?

**Figure 6:**
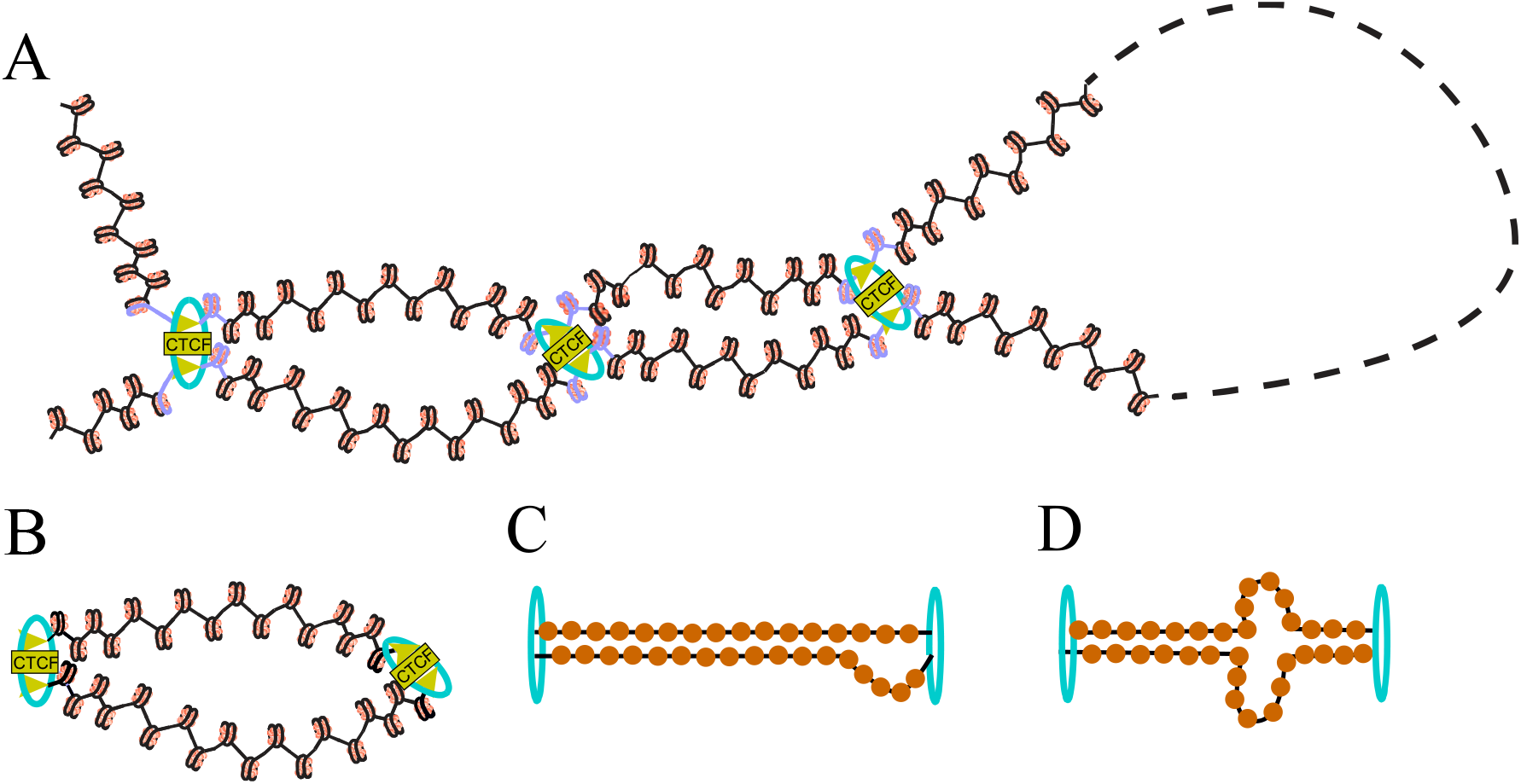
Structural homology-driven interactions in CTCF-organised nucleosome arrays. A) CTCF proteins (yellow boxes) together with Cohesin (light blue rings) separate chromatin in thousands of isolated domains that have limited interactions with each other. Four such domains separated by three CTCF-Cohesin boundaries are shown. B) Due to topological isolation of such domains, it is possible to consider one domain independent of the surrounding chromatin. C) and D) Nucleosome arrangement inside such domain is very regular, since CTCF acts as a boundary. If the phases of nucleosome arrangement are shifted by a half-nucleosome between the two nucleosome arrays, the nucleosome arrays exhibit reduced repulsion. In this case, instead of random self-avoiding loops, nucleosome arrays may form stem-like structures if under osmotic stress.

The first question is answered in our calculations shown in Figure 6A. We have compared the potential well for a highly ordered nucleosome array (decreased fuzziness, *δ* = 1 nm) with a less ordered system (increased fuzziness, *δ* = 6.8 nm). This calculation shows that this increased ordering of nucleosome array significantly deepens and heightens the energy wells and peaks. Additionally we see an increased periodicity in the peaks, which decay over a much longer length, indicating an increase in the correlation length. (Figure 7A). Given such increased recognition energy, ordered chromatin arrays near CTCF may have the tendency to form stem structures rather than loops (Figure 6C and D). This effect may be facilitated by the general osmotic stress and confinement pressure in the cell nucleus, as well as the additional pressure of DNA supercoiling acting near CTCF sites due to Cohesin molecular motors, which actively form chromatin loops at CTCF sites^54^.

**Figure 7:**
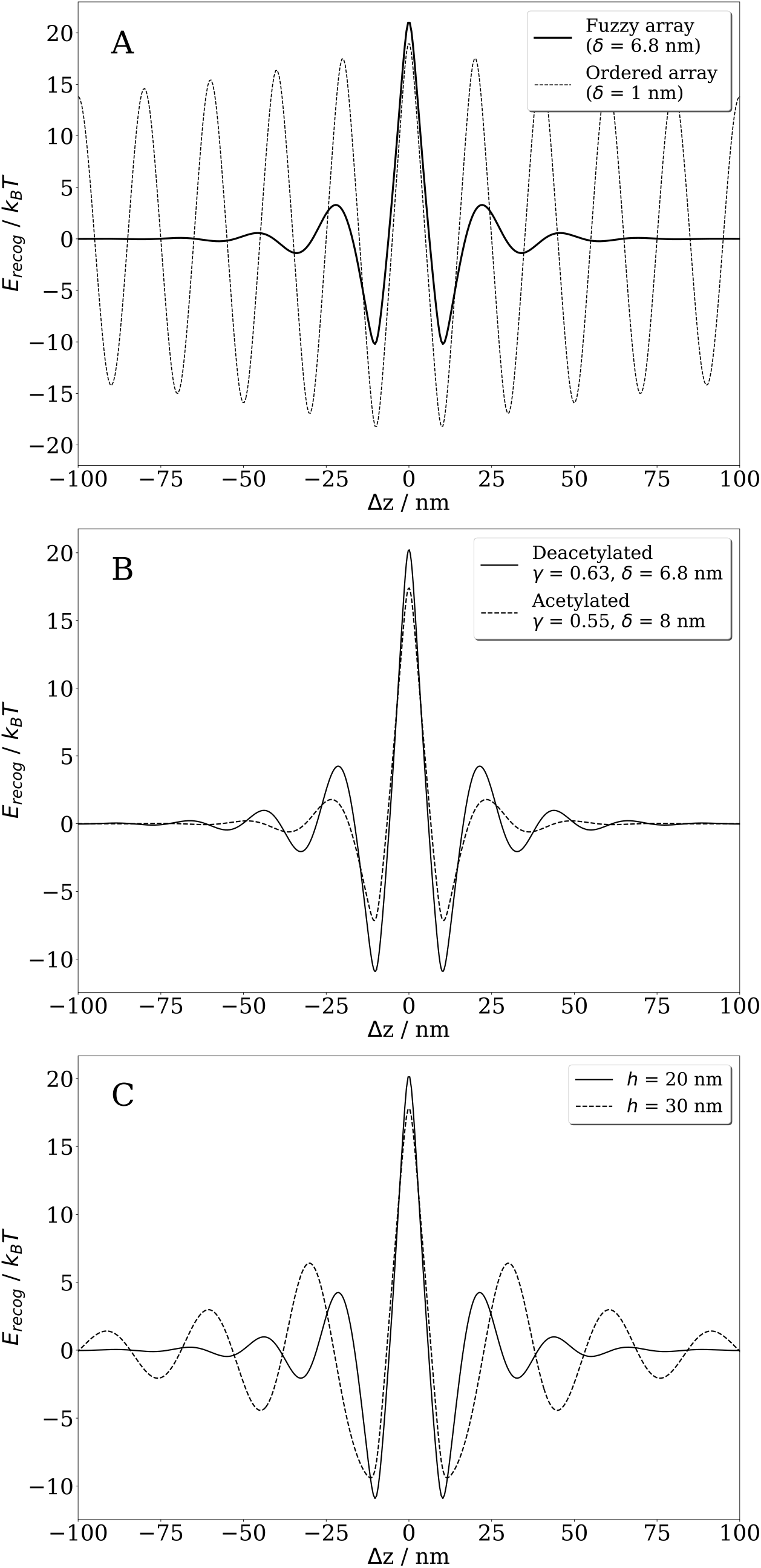
Biological effects of structural homology of nucleosome arrays. A) Ordered nucleosome arrays near CTCF (δ = 1 nm) are characterised by significantly more structured recognition potential well than “fuzzy” nucleosome arrays (δ = 6.8 nm). B) Histone acetylation modelled by the change in electrostatic potential leads to a decreased depth of the potential well. Deacetylated nucleosome array: γ = 0.63, δ = 6.8 nm; acetylated nucleosome array: γ = 0.55, δ = 8 nm. C) The increase of NRL leads to the widening of the recognition well (average distance between nucleosomes, h = 20 nm and h = 30 nm).

### Histone acetylation can decrease attraction of nucleosome arrays, but facilitate their sliding

Histone modifications are essential elements of gene regulation, catalysed by a number of specialised enzymes^55^. We therefore expect that many histone modifications will affect the structural homology recognition of nucleosome arrays. Let us consider, for example, histone acetylation. When a lysine residue on the histone tail is acetylated, a negative charge is added, which decreases the overall positive histone charge, making the nucleosomes significantly less stable^56,57^. Since acetylation destabilises the nucleosomes, it also destabilises the precision of their genomic locations, and hence increases the fluctuations in nucleosome positioning. Acetylated nucleosomes are usually found in active, less compact chromatin regions, including enhancers and promoters. In the model presented above, acetylation can be accounted for by (i) the nucleosomal charge (through *γ*), a dimensionless parameter defining the charge compensation by the histone; and (ii) the fuzziness parameter δ. Indeed, we can model the effects of acetylation and destabilisation of the nucleosome by decreasing *γ* and increasing the nucleosome position variability by increasing δ, and then exploring how this would affect the recognition well.

The combination of these two effects, as seen in Figure 7B, significantly reduces the magnitude of the recognition well. It’s worth noting, that for the set of parameters in Figure 7B, the depth of the well is still high enough to be considered as a significant trap at ∼7-8 kBT, while the energy barriers to reach this well have a significant reduction in height. Thus, acetylated fibers may slide more easily against each other, as there are smaller and fewer barriers to this paired state. This is consistent with the overall more active and less condensed nature of acetylated chromatin regions.

### NRL increase weakens attraction between nucleosome arrays

A number of studies have indicated that a change in the nucleosome repeat length (NRL) regulates the functional state of the chromatin region^58,59^. In particular, the transcriptionally active euchromatin has a smaller NRL, while transcriptionally repressed heterochromatin has a longer NRL^60–63^. This is somewhat counterintuitive, as one may expect smaller distances between nucleosomes in the chain when they are tightly compacted. However, this larger spacing may also facilitate the condensation of the nucleosome chain due to more flexibility for the bending of the stiff linker DNA between nucleosomes. The NRL also captures the (local) composition of the chromatin, e.g. the abundance of linker histones H1, which is explained both by structural and electrostatic contributions^64^. The NRL modulation by local enrichments/depletion of linker histones H1 has been recently shown to play important functional roles in regulating gene expression^65,66^.

Within the theory developed here, we can crudely model the increase of NRL by increasing the average distance between nucleosomes, *h*, and observing how this affects the recognition well. Figure 7C shows that increasing *h* decreases the depth of the primary wells but deepens the subsidiary wells. This is accompanied by an increase in both the energy barriers and coherence length, and consequently an increase in the distance over which the interaction persists. This indicates that, regardless of the homology of the sequences, they will still interact at larger shifts and can associate more easily, as required in the formation of the more condensed structure of heterochromatin. Additionally, the increase in barrier height coupled with the reduction in the energy well depth indicates a reduction in the ability of the fibers to align at higher NRLs.

The mathematical model presented here considers a parallel juxtaposition. However, it is rather obvious that such juxtaposition is ideal, and while it is possible for external factors to provide it, in reality the fibres will not be ideally parallel along their entire length. Chromatin states are characterised by regions with local alignment of nucleosome arrays; this alignment is never ideal, but the arrays do not necessarily need to be parallel to observe these attraction effects. Earlier analysis performed for bare DNA has shown that undulations of DNA molecules in nearly parallel juxtaposition can only enhance the structural effects of DNA-DNA interactions^67^.

It is worth noting, that the heterochromatin/euchromatin distinction includes both differences in NRL and differences in histone modifications (as well as differences in the associated proteins, etc). As we have shown above, the effects of these components are not always acting in the same direction with respect to the recognition of nucleosome arrays. For instance, both longer NRLs associated with heterochromatin and histone acetylation associated with euchromatin weaken the attraction of nucleosome arrays. Thus, the combined effect of these different components is defined by a subtle interplay of many contributions.

Note also that the latter effect of the weakening of nucleosome array attraction at larger NRLs is also relevant in the case of nucleosome arrays bounded by CTCF depicted in Figure 6A. Indeed, since nucleosomes closest to CTCF have smaller NRLs^51^, the effect of nucleosome array attraction will weaken further away from CTCF, due both to the loss of array ordering (Figure 7A) and also the NRL increase (Figure 7C). Thus, CTCF sites provide ‘focal points’, around which the nucleosome arrays interact the most.

### Effect of structural homology of nucleosome arrays on homologous DNA recombination

Let us now consider the effect of structural homology of nucleosome arrays on the process of homologous DNA recombination. Structural homology arises due to symmetric patterns of nucleosome arrangements between two chromatin fibres along the genomic coordinate. As justified earlier, this symmetry may involve the homology of the underlying DNA sequence, and so we can link the structural homology recognition well to the effect of homologous recombination, and the alignment of genes.

Understanding the ‘energetics’ of recognition itself does not in itself provide much information on how these molecules can reach this position of full juxtaposition *in vivo*. Nevertheless, determining the ‘recognition potential well’ as a function of axial shift is a net step that rationalizes a possible driving force for homologue-to-homologue juxtaposition, in a hypothetical homology search process of two DNA molecules sliding against each other, before they recognise the match.

The comparison of the recognition well obtained here against the helix-specific recognition well between bare DNA molecules at these length-scales (Ref. 4), shows that this structural homology recognition is not negligible. We can speculate on a two-step mechanism by which DNA homology recognition can occur in chromatin: 1) the fibers in their 10-nm form slide along each other until they reach near-alignment, at which point 2) the nucleosomes within the juxtaposition window can be moved/removed to allow for full alignment of the DNA section according to helix-specific interactions.

It is important to note that an ‘attractive’ minimum in the recognition energy potential well does not necessarily indicate direct electrostatic attraction between fibers. By analysing the interaction between chromatin fragments, we see that they repel each other along their entire length, and the minima are regions of reduced repulsion near full juxtaposition. When in confinement, the fibers will adopt configurations with the least repulsion, supporting the idea that confinement potentially plays a large role in the homology search process (see *Conclusion*).

In the search for homology, if there are large barriers between the wells, it may make sense for the 10-nm fibers to detach from each other, i.e. jump up out of close proximity, and then land at the next favourable, axially shifted juxtaposition^42^. This assumes that the fibres follow a ‘pure sliding process’, involving the simple movement of one rigid fibre over another, as described by our model. However, the 1D barriers present here can be large, which can obstruct this simple parallel sliding. For long nucleosome arrays, a reptation mechanism of sliding may be preferred – a process that may be associated with its own free energy activation barriers, not calculated here. It is entirely possible that these ‘reptation energy barriers’ are lower than the 1D barriers that emerge here, facilitating the sliding process. There are a number of indications that homologous sequences can be spatially co-located, e.g. during DNA damage repair^68,69^. Additionally, chromosome mobility increases during repair processes^70^, which facilitates bringing homologous regions closer together. We will not go into this in the present paper, but we wish to keep in mind that we must not take the 1D-barriers literally, as the relative motion of DNA in the homology search could proceed on a multi-dimensional potential energy surface, which will require separate random-dynamics analysis.

There does exists experimental evidence of an inverse relationship between the nucleosome density and the efficiency of homologous recombination^31–33^. While the likely effect of an increased nucleosome density is to restrict required enzymes from accessing the bare DNA, there may potentially be a more fundamental reasoning behind this observation which does not rely on enzymes. From the theory presented here, we can see that this increased density (by reducing the nucleosome array fuzziness, *δ*) both deepens the potential wells and raises the energy barriers to alignment. This increases the difficulty for these fibers to come into near-full juxtaposition, requiring a much higher ‘jump & land’ as previously described, hence leading to the observed relationship in Refs. 31-33. The effect of reducing the nucleosome density significantly lowers these barriers, facilitating the path to near full juxtaposition. The two-step mechanism speculated above is therefore still valid as proteins and enzymes required for the next steps of homologous recombination are free to access the relevant nucleosome-depleted regions of the chromatin fibers once they are in full alignment as a result of nucleosome sliding and the helix specific interactions between bare DNAs.

## IV. CONCLUSIONS

In this work, we have developed the theory of structural homology recognition by nucleosome arrays and have applied it to several biological scenarios observed *in vivo*, which can be briefly summarised as follows:

1. Figure 7A shows that more ordered homologous nucleosome arrays can have a stronger attractive component of interaction than less ordered ones. Increased ordering can be provided by DNA sequence repeats, nucleosome-disfavouring DNA sequences or strongly bound proteins such as CTCF (Figure 6A).
2. In such arrays, nucleosomes closer to CTCF are more ordered and have smaller NRLs, which means they act as foci of interactions between chromatin fibers.
3. Sections of nucleosome arrays between boundaries may form stem-like structures rather than free loops, contrary to current assumptions in the field (Figures 5C and D).
4. Histone acetylation weakens structural homology recognition between nucleosome arrays but facilitates chromatin fiber sliding with respect to each other.
5. Chromatin regions with larger NRLs are characterised by weaker structural homology recognition.
6. Structural homology recognition may facilitate homologous recombination, detailed below.

With respect to the process of homologous recombination, the helix-specific DNA recognition is usually expected to be important after DNA is stripped of histones. However, the analysis in this paper shows that an earlier opportunity may be utilised for the pairing interaction in chromatin, arising from the sequence-specific distribution of nucleosomes along the DNA. This results in a noticeable ‘*recognition well’* which allows close alignment of homologous DNA sequences simply as a result of the similar nucleosome positioning on homologous sequences. Full juxtaposition of the DNA could then be achieved in a follow-up stage by exposure of the bare helical structure through sliding or the stripping of nucleosomes.

Currently our model allows only for rigid chromatin fibers, where the nucleosomes are fixed in position. Work has already been done to model torsional softness for DNA, by adding an elasticity Hamiltonian to the interaction Hamiltonian^47^. Using a similar approach, it will be possible to include these elements of flexibility in chromatin. Additionally, this theory assumes that the nucleosome arrays are straight and parallel within the juxtaposition window. As for the elasticity, we can pull on earlier work to include tilted interactions where the fibers may not necessarily be parallel to each other^71^, as well as a flexible juxtaposition between the two fibers^67^. This will aid in broadening our picture of the recognition energy landscape. However, before any further development of the theory, it would be most valuable to perform single molecule experiments involving the sliding of chromatin fibers against each other. The results of such in vitro experiments would aid the verification of the results of this model. Indeed, the model on which our analysis was based is obviously crude. The systematic experimental verification of the conclusions of this analysis in a test tube (and computer simulations), would be needed.

Furthermore, nucleosome positioning is a dynamic process in the cell; they are not static, but rather fluctuate about their positions in time. Our present study constitutes a time averaged approach to the problem. Hence, an area of further interest would be to see how the discussed effects will reveal themselves in the recognition dynamics^42^. In doing this, we can develop simulations using Langevin or Brownian dynamics to gain a better picture of the system.

The model does not paint a picture of the environment in which the chromatin exists in the cell nucleus. The first step in that direction would be extending the current picture by taking into account chromatin confinement.

Still, one of the most interesting results of this investigation is the demonstration of a strong effect of different parameters of the model on the ability of homologous recognition of the chromatin fibers. By understanding this influence, realistic parameter alterations can be suggested to aid this important precursory process in DNA repair and meiosis. As a long shot, this could unravel new routes that may help minimize mistakes in recombination and thus aid the avoidance of diseases that can result from such errors.

## AUTHOR CONTRIBUTIONS

AAK conceived the work, AAK and JGH formulated the model and developed the theory in consultations with VBT, JGH performed all calculations and their preliminary analysis, all authors examined the results, with VBT focused on the analysis of the biological implementation of the model. All authors wrote the paper.

## ACKNOWLEDGEMENTS

JGH thanks Imperial College London for funding of a President’s PhD Scholarship. VBT was partially supported by the Wellcome Trust grant 200733/Z/16/Z. We are thankful to Daniel Jost (Laboratory of Biology and Modeling of the Cell, CNRS, ENS Lyon.) for useful discussions.

## V. APPENDIX

### 1. Recognition Well for Sliding Genes – Derivation of Eq. (6) of the main text

Using Eqs.(2) –(4) of the main text, we first modify the interaction Hamiltonian to include the contributions from both the nucleosomes and the DNA. Following through the derivation of the interaction Hamiltonian using this charge density, we obtain three terms:

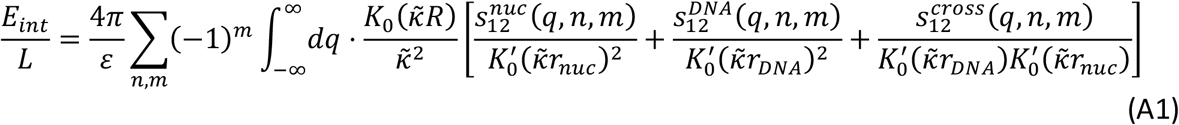

Where 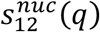 and 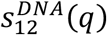 are the charge density correlation functions for the nucleosomes and DNA charge distributions respectively, and 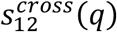 contains the cross-terms between nucleosomes and DNA. All other parameters are defined in the main text. When we consider the recognition well, we must consider the difference between homologous and non-homologous interaction energies, i.e. *E*_*recog*_ = *E*_*hom*_ − *E*_*nonhom*_. Examining Eq. (A1), we find that the only term that differs between the homologous and nonhomologous cases, in this model (which is not the case for KL-theory) is the nucleosome term. Hence, we can write the recognition energy per unit length as:

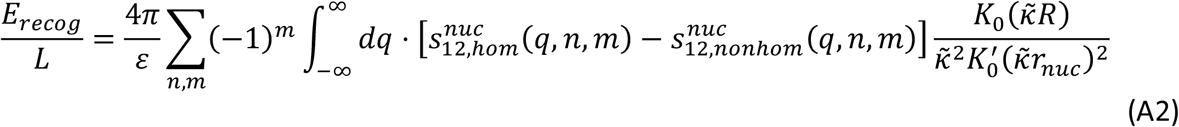

The real space surface charge densities for the nucleosomes and the DNA are given in the main text. Calculating their Fourier transforms, 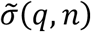, we obtain:

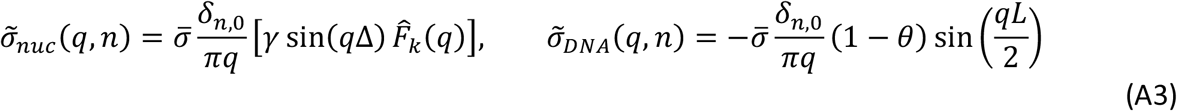

where *δ*_*ij*_ is the Kronecker delta. Again, the expression for 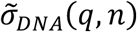, is the simplification which ignores the central point of the KL theory – the helicity of DNA which, as we have stressed, we can afford as the radius of histones is substantially larger than the DNA radius. The distribution of nucleosomes is defined by the nucleosome centres, *z*_*k*_, in:

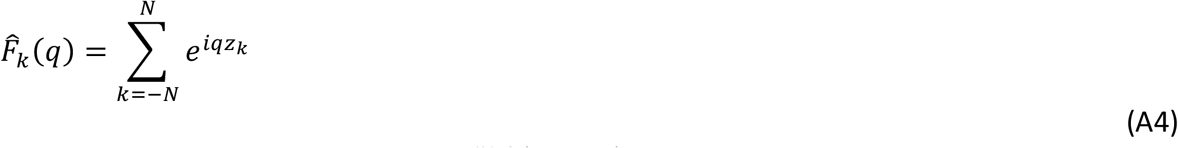

To find the recognition energy, for 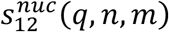, defined as in Eq. (2) of the main text, we write down the homologous and nonhomologous forms. We also now introduce the axial shift, Δ*z*, by defining 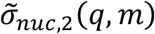 in its own coordinate system, shifted by Δ*z*. Hence, we can write 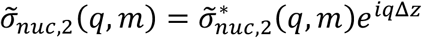. Combining this with Eqs. (A3) and (2), we find 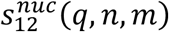:

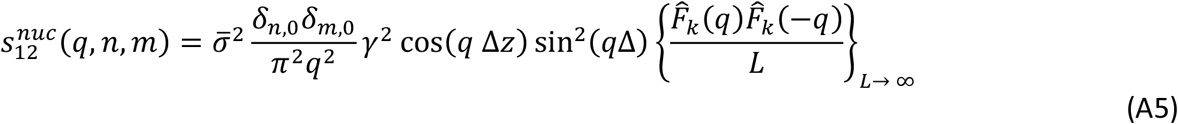

Using Eq. (6) of the main text for the nucleosome distribution, we can break down 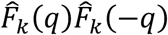 into several terms:

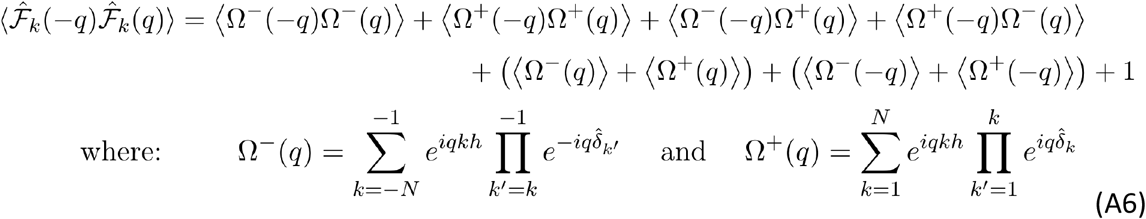

where ⟨… ⟩ denotes a statistical average. Taking the difference between homologous and nonhomologous expressions, and using ⟨Ω^+^(−*q*)Ω^+^(*q*)⟩ = ⟨Ω^—^(−*q*)Ω^−^(*q*)⟩, only two terms from (A8) remain, simplifying the calculation:

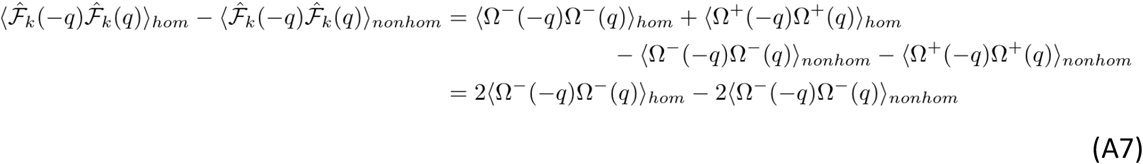

Now for the homologous case, we assume that the deviation of each nucleosome is the same for each index. Hence, the assumption we make is 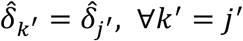. This means that the nucleosomes deviate identically along the length of each fiber, rather than having specific homologous regions. As for DNA, we assume that the deviations obey Gaussian statistics^3,40,72^. This simplification is based on results that show a non-ideal double helix can be modelled as steps with a height of one helical rise (3.4 Å), but with a non-constant twist angle, deviating from the average twist angle by ≈ 0.1 radians^73–75^. Using this, and averaging over realisations of the random deviations, we can rewrite the sum as:

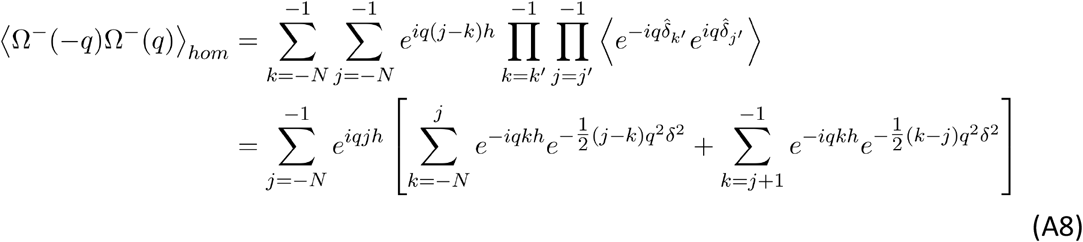

where 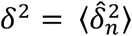. This approach is similar to that taken by in Ref. 25 used for modelling protein-DNA interactions, altering the summation ranges as required. Careful calculation of the double sums, and using *L* = (2*N* + 1)*h* reveals a lengthy expression for this average:

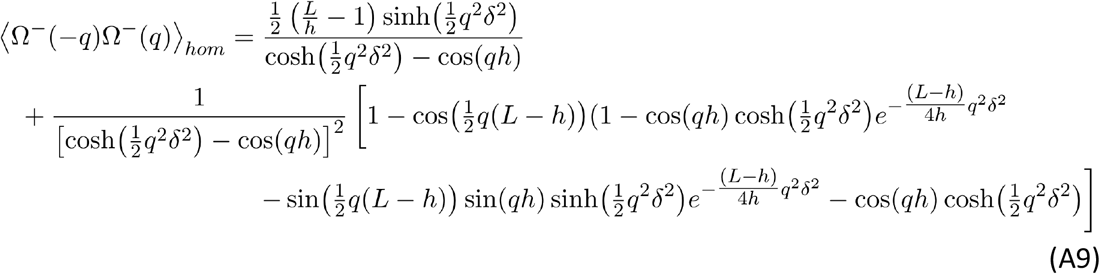

Between completely non-homologous molecules, 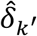’ and 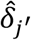’ are uncorrelated. After Gaussian averaging over the random deviations, we can write the sum as:

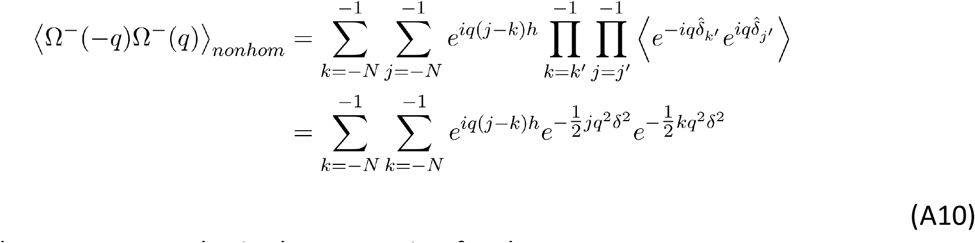

Calculating these sums, we obtain the expression for the average:

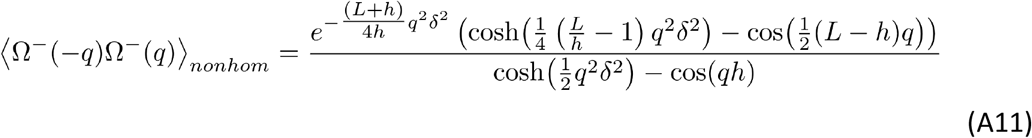

Combining equations (A2), (A5), (A9) and (A11), and calculating the recognition energy over a juxtaposition length *L*_*J*_, we arrive at Eq.(7) of the main text.

### 2. Interaction between Chromatin Fragments

For this calculation, we define the charge density correlation function as 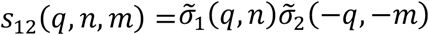 to allow for finite sized fragments. By using this, as well as shifting 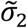 by Δ*z*, we can rewrite the interaction Hamiltonian in Eq. (A3) as:

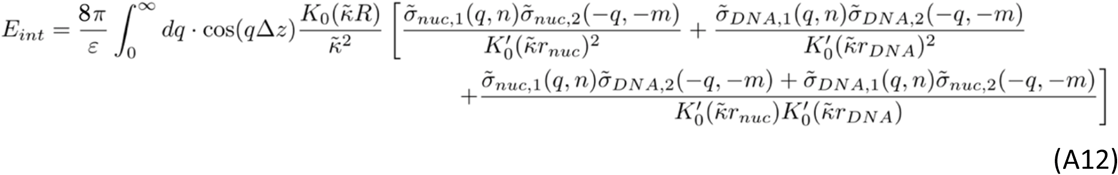

The first term in the brackets in Eq. (A12) is handled in the same way as before, except for that we now need to consider all terms in Eq. (A6). Calculating the sums and the averages, we find for ⟨Ω^±^(*q*)⟩:

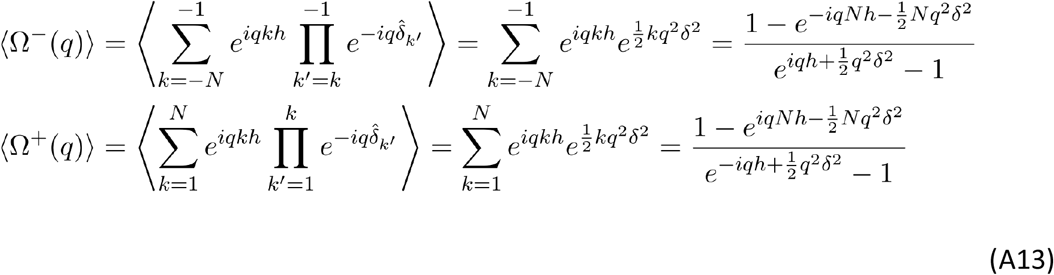

From these expressions, it is clear that ⟨Ω^−^(*q*)⟩ = ⟨Ω^+^(−*q*)⟩. Hence, the four relevant terms in Eq. (A6) can be simplified to 2(⟨Ω^+^(−*q*)⟩ + ⟨Ω^+^(*q*)⟩). Finding this expression, then expressing the complex exponentials into trigonometric functions, we obtain:

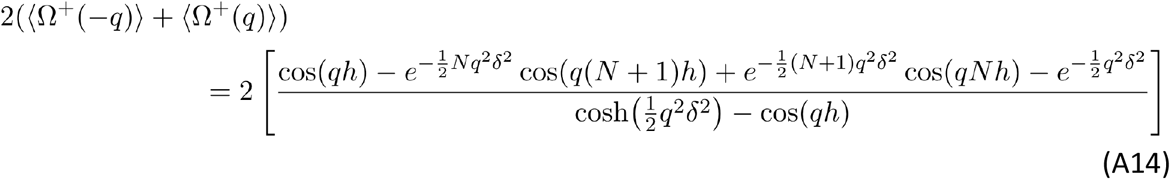

Next, we need to calculate ⟨Ω^−^(−*q*)Ω^+^(*q*)⟩ and ⟨Ω^+^(−*q*)Ω^−^(*q*)⟩. As each average contains sums corresponding to deviations on opposite ends of the fibers (either *k* > 0 or *k* < 0), there cannot be a ‘homologous’ formulation of this expression, and therefore we take the deviations to be completely uncorrelated:

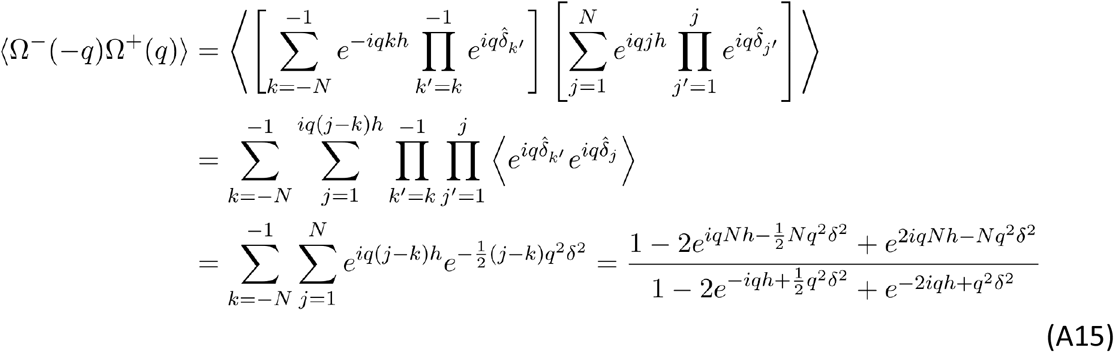

A similar procedure for the calculation of ⟨Ω^+^(−*q*)Ω^−^(*q*)⟩ gives the complex conjugate of ⟨Ω^−^(−*q*)Ω^+^(*q*)⟩. Carefully adding the terms, and converting the complex exponentials to trigonometric functions, we obtain a rather cumbersome expression for these terms:

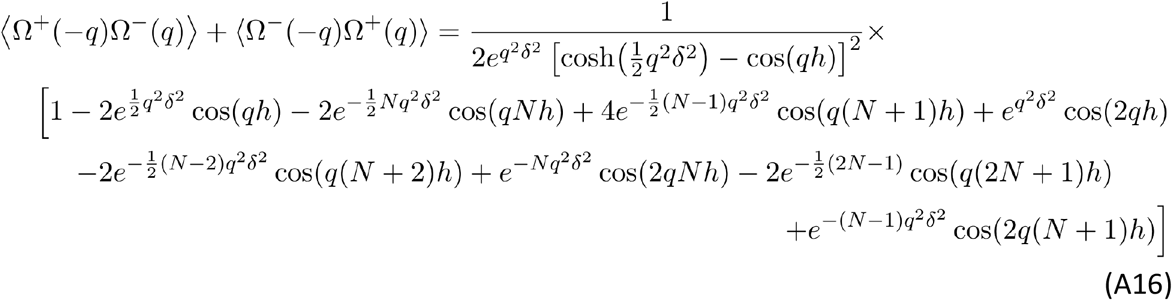

This completes the calculation of all the terms that contribute to 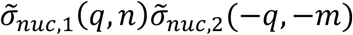. The next term to consider is the DNA interaction term. The calculation of 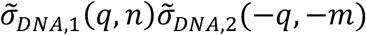 is trivial in comparison. Using Eq. (A3), we find:

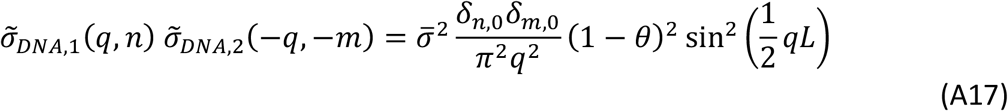

The final term to calculate is the cross term. From Eq. (A3), we see that this is equal to:

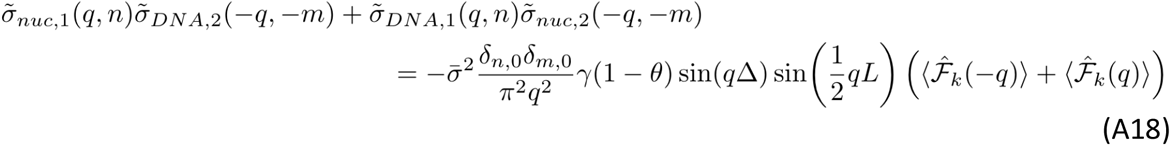

Using the definitions in Eq. (A6), we can break down the calculation of 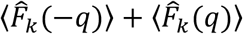 into a sum of terms that have already been calculated:

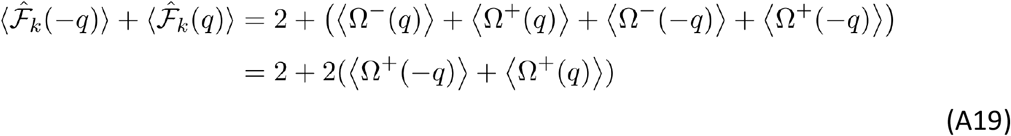

Using Eq. (A12) by combining equations (A6), (A9), (A11) and (A13)-(A19), and substituting in *L* = (2*N* + 1)*h*, we obtain the full expressions for the interaction energies of homologous and non-homologous molecule fragments of length.

Homologous:

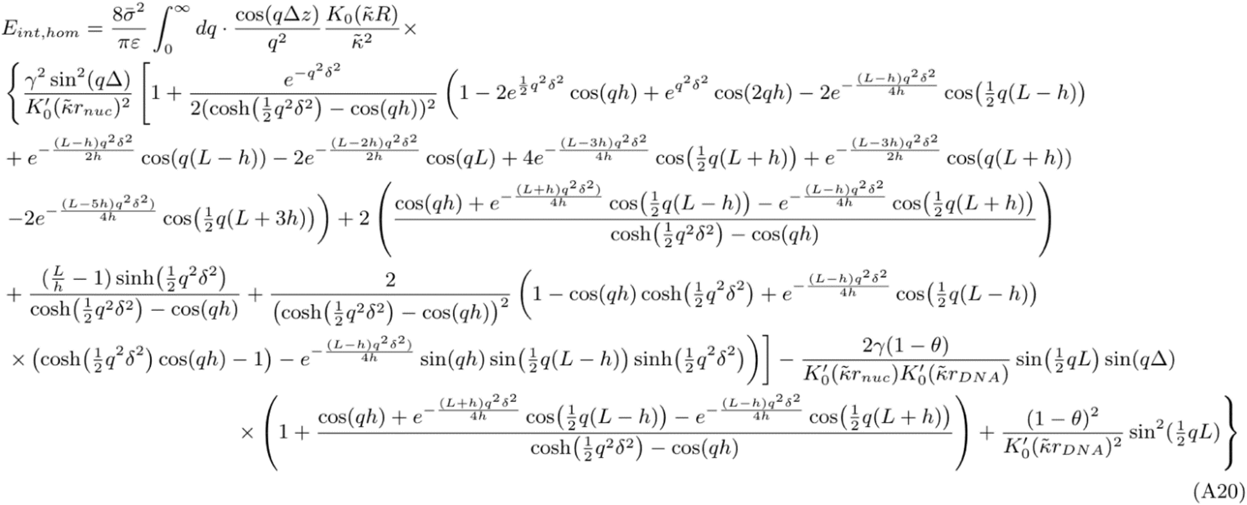

Non-homologous:

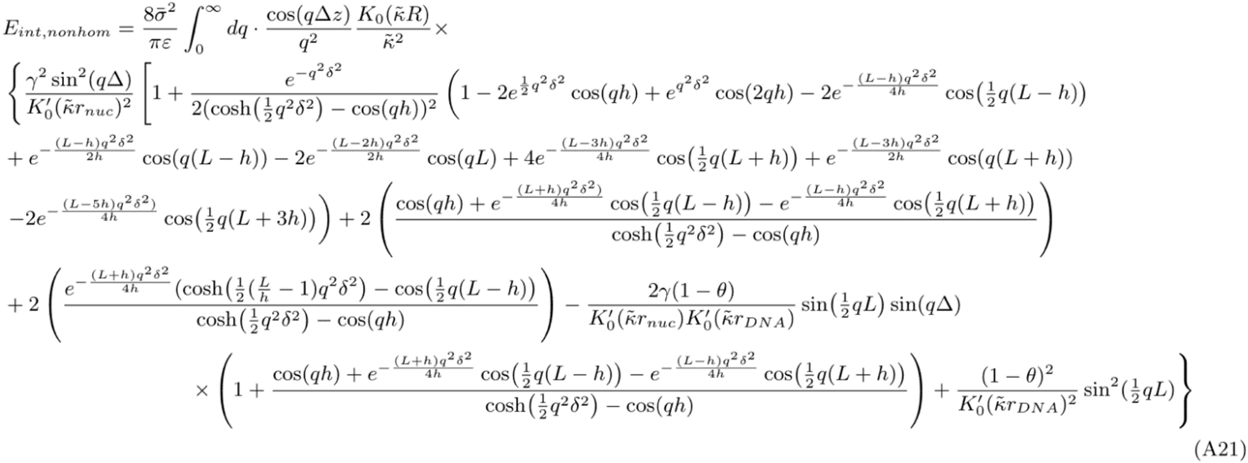

Subtracting (A21) from (A20), we obtain the form of the recognition well for chromatin fragments as Eq.(8) of the main text.

